# Early changes of ER-mitochondrial interaction in the liver of high-fat diet-fed mice

**DOI:** 10.64898/2026.04.11.717879

**Authors:** Justyna Malecka, Gabriela Chrostek, Claudio Casali, Emanuela Pessolano, Elena Mantovani, Nausicaa Clemente, Simone Reano, Giulia Pinton, Marco Biggiogera, Laura Tapella, Fausto Chiazza, Giulia Dematteis, Dmitry Lim

## Abstract

IP3R-Grp75-VDAC1 protein complex at the mitochondria-ER contact sites (MERCS) is involved in response to nutrients and control of glucose and energy metabolism, however, early alterations of the complex and MERCS in response to increased fat intake remain inconclusive. We investigated early effects of high-fat diet (HFD) on IP3R-Grp75-VDAC1 protein expression in correlation with ER-mitochondrial interaction in the liver of mice. Five-week-old mice were fed an HFD or a standard diet (SD) for 2 weeks (2W) or 8 weeks (8W). MERCS fractionation by a gradient ultracentrifugation, Western blot, transmission electron microscopy (TEM), Oroboros high-resolution respirometry were used to analyse liver tissues, while real-time PCR was used to profile genes responsive to HFD. No macroscopic morphological or functional alterations were observed in mice at 2W, while, expectedly, at 8W of HFD mice gained weight and glucose intolerance. Total IP3R protein was reduced at both 2W and 8W points by a post-transcriptional mechanism, while in MERCS, IP3R, VDAC1 and Grp75 were reduced at 8W time-point. TEM analysis revealed a significant reduction of mitochondrial coverage by MERCS, mitochondrial fragmentation and shortening of ER-mitochondria distance already at 2W time-point. Mitochondrial function and metabolism were largely spared. Markers of altered protein homeostasis such as Lmp2, Mecl-1 and Lmp7 showed an early upregulation. In conclusion, HFD induces early alterations in liver MERCS that precede gain of weight and glucose intolerance, suggesting their primary role in obesity and metabolic diseases and as potential therapeutic target.

## INTRODUCTION

Hypercaloric and/or hyper-processed fat-rich food intake (high-fat diet, HFD), causing metabolic dysregulation and obesity with associated complications such as non-alcoholic fatty liver disease (NAFLD) and non-alcoholic steatohepatitis (NASH), represent a global healthcare challenge [1,2]. These disorders are also associated with insulin resistance, type II diabetes (T2D) and represent a risk factor for cardiovascular diseases and age-related dementias, such as Alzheimer’s disease [3–7]. Although cellular and molecular mechanisms of these diseases are being investigated, growing number of reports point to the dysfunctions of the endoplasmic reticulum (ER) and mitochondria, and, specifically, on the mis-communication between these two organelles at the so-called mitochondria-ER contact sites (MERCS) [8–11]. MERCS, formed by the juxtaposed outer mitochondrial membrane (OMM) and mitochondria-associated ER membrane (MAM), host numerous proteins involved in a number of key cellular processes such as ER-mitochondrial transfer of lipid and Ca^2+^, mitochondrial bioenergetics, RedOx balance, protein synthesis and autophagy as well as in the response of cell to stress in forms of apoptosis and ER-stress/unfolded protein response (UPR) [12–17]. A low-affinity but high capacity mitochondrial Ca^2+^ uptake, which drives energetic metabolism through Ca^2+^ activation of mitochondrial enzymes, occurs in the mitochondria located closely to the ER through MERCS [18,19]. Dysregulation of the ER-mitochondrial Ca^2+^ flux has been associated with HFD-induced obesity and insulin resistance in adipose tissue, liver and muscle [8–10,20]. The protein complex, implicated in Ca^2+^ transfer is composed of Inositol-1,4,5-trisphosphate receptors (IP3R) on the ER membrane and voltage-dependent anion channel 1 (VDAC1) on the OMM, tethered to each other by glucose-regulated protein 75 (Grp75) [17,21]. Other IP3R-interacting proteins, implicated in the regulation of Ca^2+^ flux from the ER include non-opioid sigma 1 receptor (SIGMAR1), translocator of outer mitochondrial membrane 70 (TOM70) and parkinsonism associated deglycase DJ-1 [17,22–25]. The efficiency of ER-mitochondrial Ca^2+^ transfer depends primarily on two MERCS parameters: i) transversal distance between OMM and MAM and ii) extension of the interface of the juxtaposed OMM and MAM [14,17,26,27]. Recently it has been demonstrated that the dependence on the transversal distance (gap size) has a bell-shaped profile: Ca^2+^ flux is suppressed at the distances ≤10nm and ≥30nm and is optimal at a distance about 20nm [28]. Furthermore, more extended interface, generally, correlates with a higher Ca^2+^ flux [8,14,17]. MERCS are highly dynamic structures, though the number of contacts, their interface and gap size change according to cell condition. Although somewhat contrasting results have been published regarding the strength of the interaction between ER and mitochondria in mice with genetic and induced obesity, reports suggest that alterations of MERCS represent an early phenomenon in pathogenesis of HFD-induced metabolic dysfunction and obesity, however, the expression of components of the Ca^2+^ transferring unit in MERCS, in correlation with their physical parameters is not fully understood [10].

Recently we reported that in a cohort of mice fed with HFD for only two weeks, i.e., prior signs of obesity and metabolic alterations, a methyltransferase Enhancer of Zeste Homolog 2 (EZH2) mediated trimethylation of lysine 27 in histone H3 (H3K27me3), a major epigenetic modification towards a massive transcriptional repression [29]. Here, following European Directive 2010/63/EU, mandating the “3Rs” principle (Replacement, Reduction, Refinement), we used the same cohort of mice to investigate the expression of IP3R, Grp75 and VDAC1 in total lysates and isolated MERCS at presymptomatic stage (2 weeks, 2W) and at the stage with overt signs of obesity and metabolic dysregulation (8W) in correlation with MERCS ultrastructural changes, mitochondrial function and transcriptional markers of HFD-related dysregulation.

## RESULTS

### General characterization of HFD mouse model

Body weight did not differ between groups at baseline. After two weeks of dietary intervention, mice fed the hypercaloric diet did not display a statistically significant increase in body weight compared with control animals. In contrast, a significant increase in body weight was observed after eight weeks (8W) of HFD exposure (Table 1).

**Table 1.** HFD 8W but not 2W affects mice body weight and fasting glycemia. *** = p < 0.001 vs. time-corresponding SD. Data are presented as mean ± s.d.

|  | SD |  |  | HFD |  |  |
| --- | --- | --- | --- | --- | --- | --- |
| | Basal<br>(Mean±SD) | 2W<br>(Mean±SD) | 8W<br>(Mean±SD) | Basal<br>(Mean±SD) | 2 W<br>(Mean±SD) | 8 W<br>(Mean±SD) |
| Body weight<br>(g) | 20.75±2.5<br>5 | 22.87±2.3<br>1 | 29.15±2.52 | 20.91±3.3<br>3 | 25.52±3.40 | 39.38±5.13*** |
| Fasting glycemia<br>(mg/dL) | 104.7±14.<br>2 | 89.5±14.1<br>6 | 90.38±18.3<br>2 | 101.1±12.<br>4 | 104.6±23.3<br>5 | 146.0±11.24**<br>* |

Regarding fasting glycemia, no statistically significant differences were detected between groups after two weeks (2W) of diet administration, whereas a significant increase in glycemia was evident after eight weeks of HFD treatment (Table 1).

### Early down-regulation of total IP3R and depletion of IP3R, VDAC1 and Grp75 from MERCS at 8W of HFD

Mitochondria-ER contact sites (MERCS) fraction was prepared from liver tissue of mice according to a validated protocol [30] with modifications specified in the Methods section. Total homogenates (HOM), crude mitochondrial fraction (MIT) and MERCS (MERC) were subjected to Western blot (WB) analysis for a quality control. As shown in Figure 1, ER membrane located IP3R was present in homogenate and MERC but absent from mitochondria. Mitochondrially-located VDAC1/3 and Mfn1 as well as cytosolic Grp75 were present in all fractions, but were enriched in MERC compared with crude mitochondria. Grp75 forms a complex with IP3R and VDAC1 [21], therefore its presence in crude mitochondrial fraction was expected. Drp1, a mitochondrial fission protein located in the cytosol, is absent from mitochondria and MERCS, which could be expected because Drp1 is recruited to mitochondria only during fission process [31]. Ubiquitous GAPDH was found in all fractions, while actin, abundantly present in total homogenate, was barely visible in crude mitochondria or MERCS. Altogether, these results suggest successful isolation of MERCS from the mouse liver.

**Figure 1.**
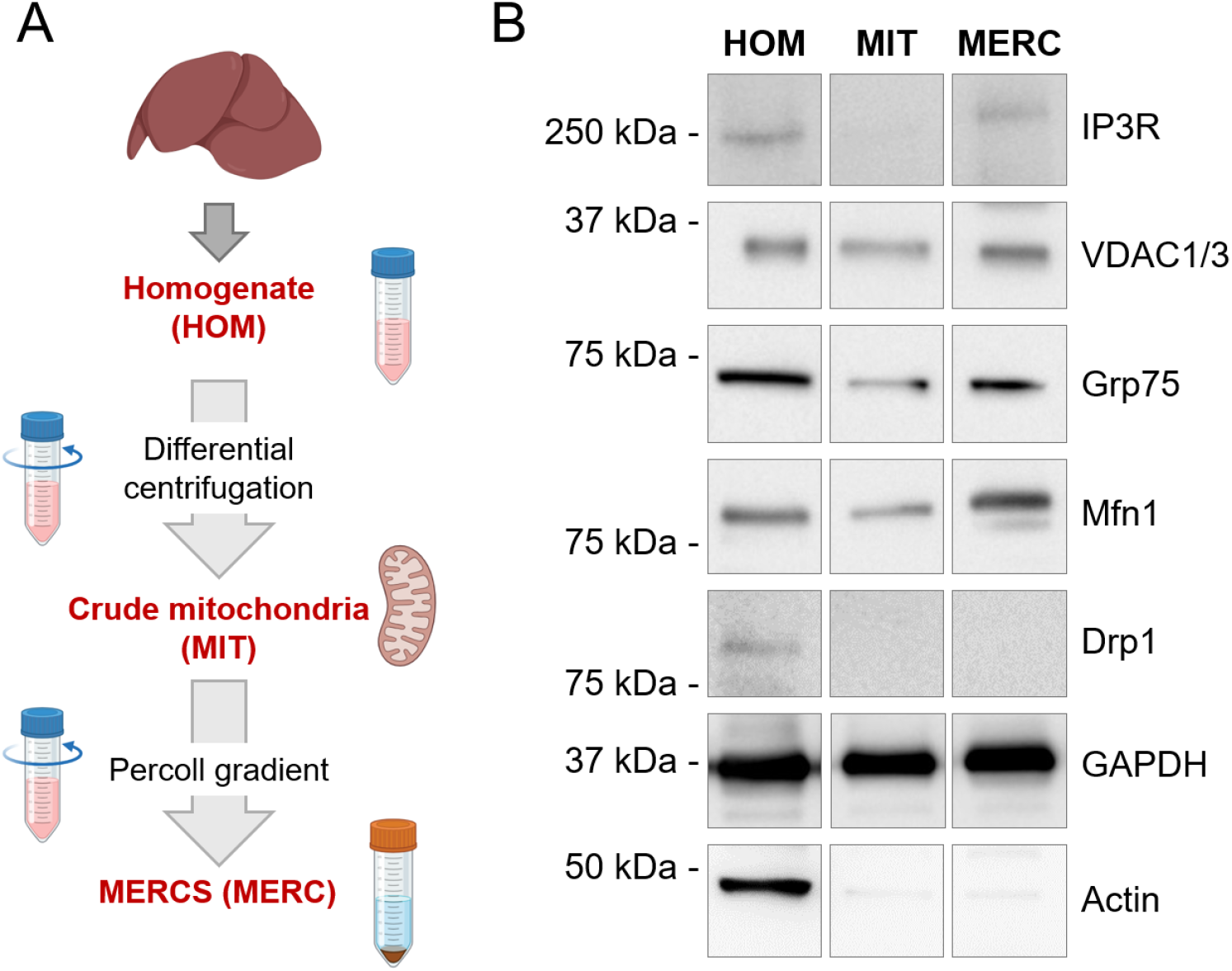
Isolation and characterization of MERCS from mouse liver. (**A**), cartoon depicting main steps in the isolation of mitochondria-ER contact sites (MERCS). (**B**), representative Western blot images of IP3R, VDAC1/3, Grp75, Mfn1, Drp1, GAPDH and Actin in total homogenates (HOM), crude mitochondrial fraction (MIT) and MERCS (MERC). All fractions were run on the same SDS-PAGE, which included other conditions, at least in triplicate. Representative bands were cut and assembled in a figure. Full-size uncut Western blot membranes are present in Supplementary materials.

Next, we analysed the expression of proteins of interest. As shown in Figure 2, IP3R was significantly reduced in total homogenates already at 2W time-point but not in MERCS, in which it was unchanged. No alterations of Grp75 or VDAC1/3 were found neither in total homogenates nor in MERCS.

**Figure 2.**
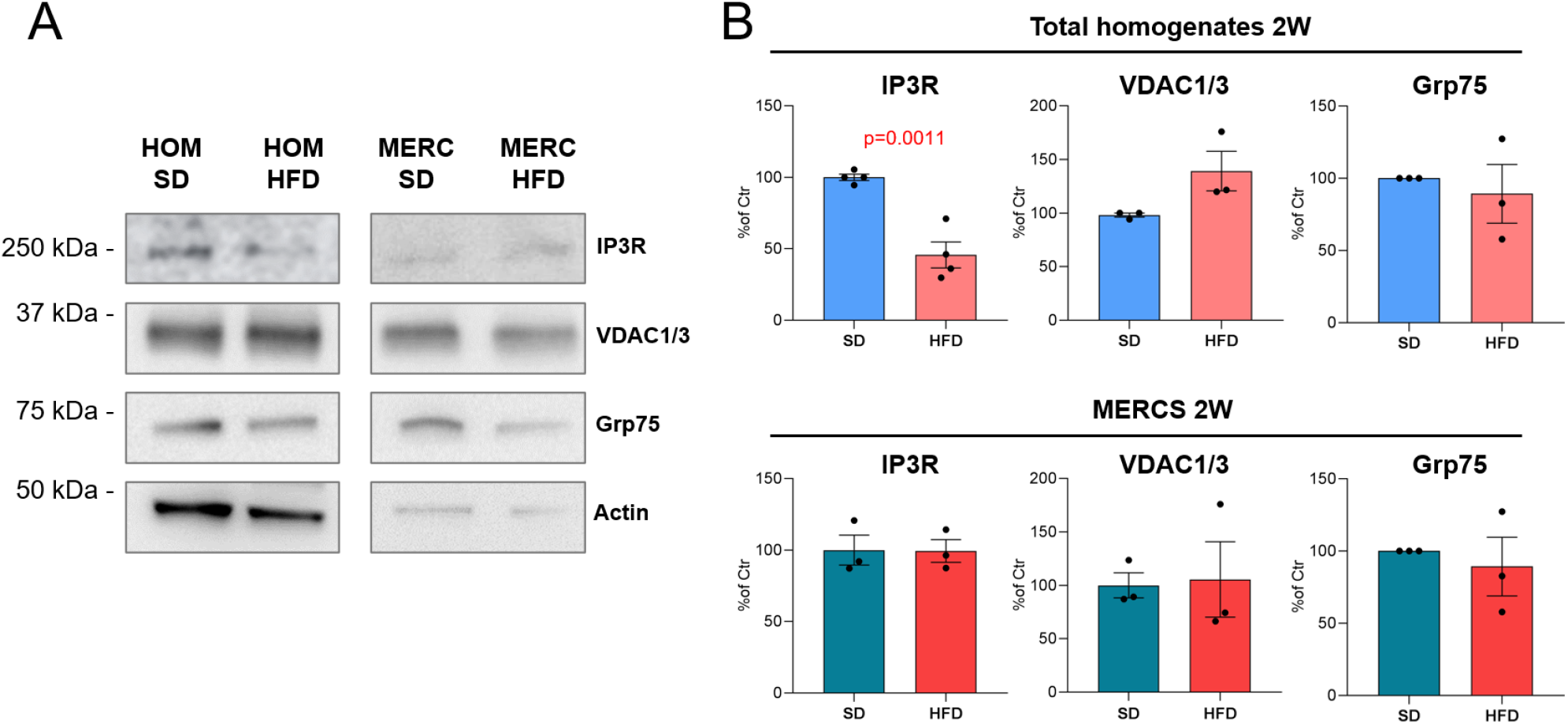
Western blot assessment of IP3R, VDAC1 and Grp75 protein expression at 2W of HFD. Representative WB images (**A**) and quantifications of IP3R, VDAC1/3 and Grp75 (**B**) in SD and HFD-fed mice at 2W time-point. In (**A**), total homogenates and MERCS were run on the same or separate SDS-PAGE, while SD and HFD conditions were always run on the same gel. In (**B**) data are expressed as mean ± s.e.m. from 3-4 independent WBs. P values are shown for p < 0.05.

At 8W time point, IP3R but not VDAC1/3 or Grp75, were downregulated in total homogenates (Figure 3). VDAC1/3 immunoreactivity somewhat showed high level of variability between samples with a tendency to downregulation. Strikingly, all proteins of interest were significantly downregulated in MERCS at 8W of HFD: IP3R and VDAC1/3 were reduce by about 50%, while Grp75 was reduced by about 80% (Figure 3). These results suggest that at a whole cell level IP3R is downregulated as early as 2W during HFD, while all studied proteins are depleted from MERCS at 8W of HFD time-point.

**Figure 3.**
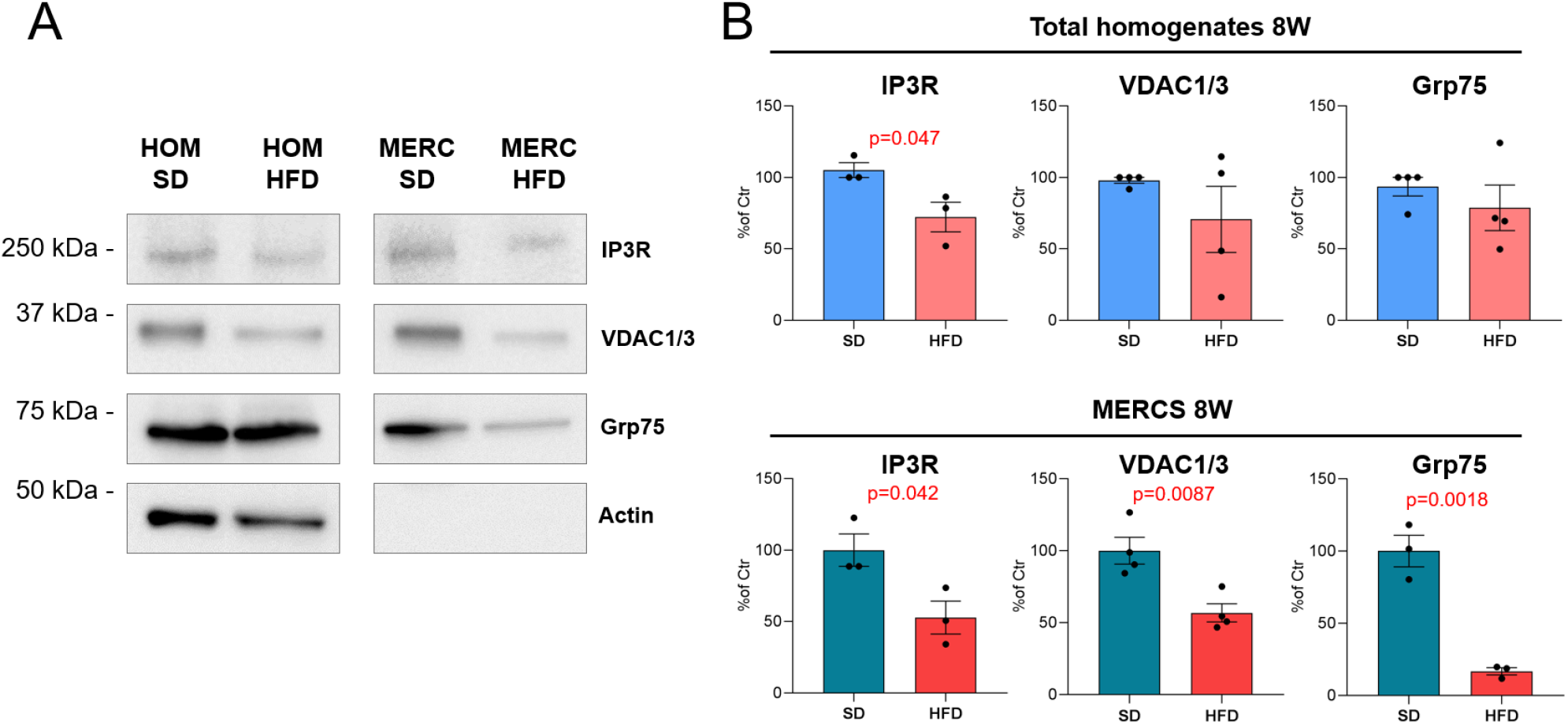
Western blot assessment of IP3R, VDAC1 and Grp75 protein expression at 8W of HFD. Representative WB images (**A**) and quantifications of IP3R, VDAC1/3 and Grp75 (**B**) in SD and HFD-fed mice at 8W time-point. In (**A**), total homogenates and MERCS were run on the same or separate SDS-PAGE, while SD and HFD conditions were always run on the same gel. In (**B**) data are expressed as mean ± s.e.m. from 3-4 independent WBs. P values are shown for p < 0.05.

To investigate whether the reduction of IP3R protein occurred at a transcriptional or post-transcriptional level, and if there were changes in mRNA expression of VDAC1 and Grp75 we performed qPCR analysis using specific oligonucleotide primers against respective mRNA species. First, we assessed levels of expression of three IP3Rs isoforms. We found that, in line with previous reports [32], IP3R2 is the most expressed isoform in liver, followed by IP3R1 isoform. IP3R3, instead was expressed at extremely low levels, and was not taken in consideration in further assessment (Figure 4A). As shown in Figure 4B, no alterations were found in mRNA expression of either IP3R1 or IP3R2 between SD and HFD at both time-points. Similarly, no difference was found in mRNA expression of VDAC1 or Grp75 between SD and HFD at both 2W and 8W time-points (Figure 4C). These results suggest that the reduction of IP3R expression in total liver homogenates and the depletion of IP3R, VDAC1 and Grp75 from MERCS at 8W of HFD occurred at a post-transcriptional level.

**Figure 4.**
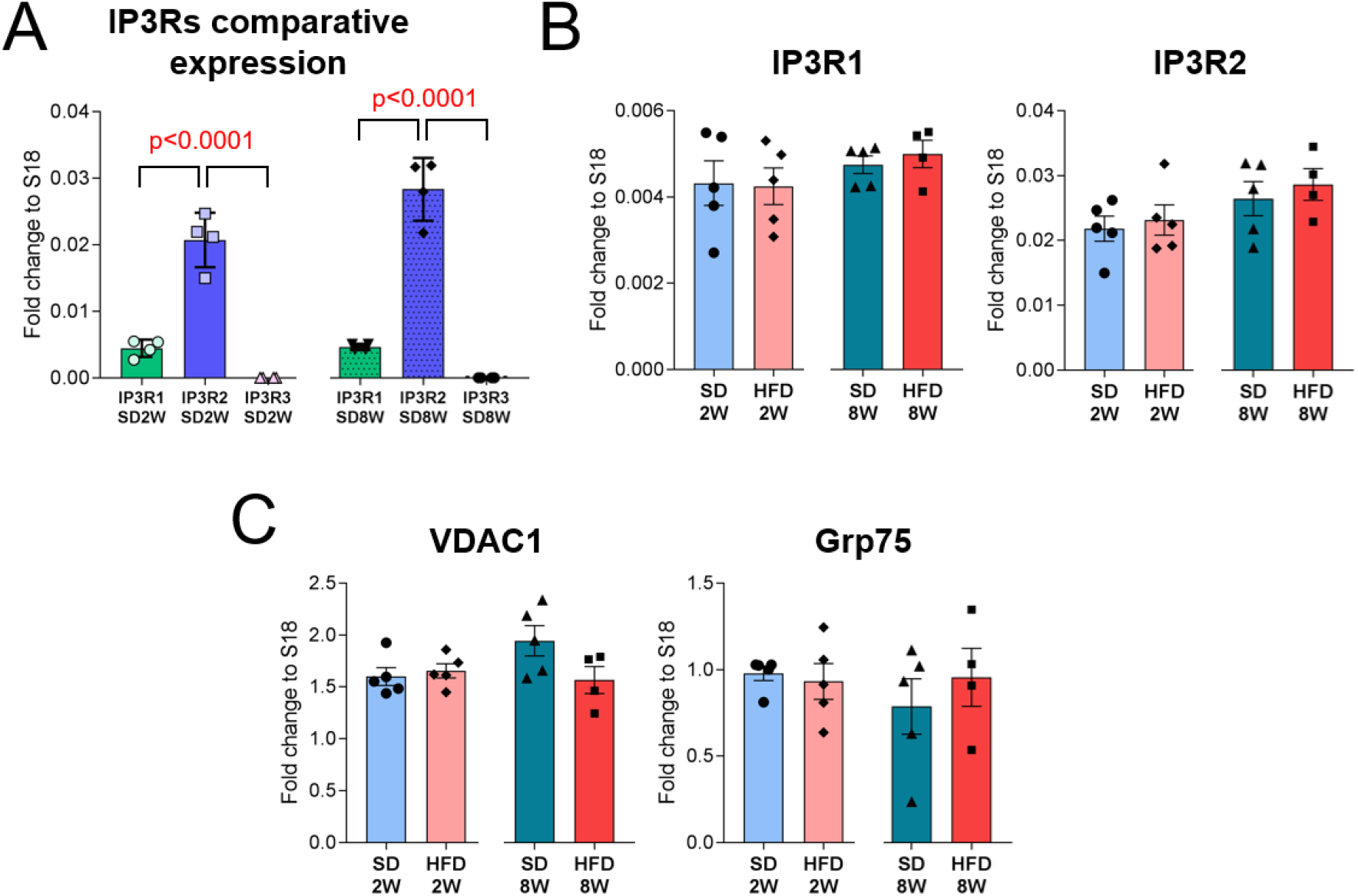
qPCR assessment of IP3R, VDAC1 and Grp75 mRNA expression. (**A**) Relative expression of IP3R isoforms in SD-fed mice at 2W and 8W. Relative expression of IP3R1 and IP3R2 (**B**) or VDAC1 and Gpr75 (**C**) in the liver of HFD-fed mice as compared with SD-fed mice. Dot plot graphs show mean ± s.e.m. of the gene expression fold change normalized to S18, each dot represent an independent mouse for n = 4-5 mice per condition. P values are shown for p < 0.05.

### Early disruption of MERCS and shortening of ER-mitochondrial distance in the liver of HFD-fed mice

To investigate whether the expression of proteins correlated with ultrastructural changes of MERCS, we performed transmission electron microscopic (TEM) analysis of liver tissue of mice fed with HFD for 2W or 8W. An overview of hepatocyte’s ultrastructure from SD-fed mice at 7500x magnification shows abundant presence of mitochondria and delimited, round, electronically clear droplets, identified as lipid droplets (Figure 5, SD 2W and SD 8W). Analysis of HFD-fed mice at 2W time-point shows regular ultrastructure with abundant lipid droplets (Figure 5, HFD 2W). Analysis of HFD-fed mice at 8W time-point showed presence of enlarged lipid droplets, and steatosis. (Figure 5, HFD 8W).

**Figure 5.**
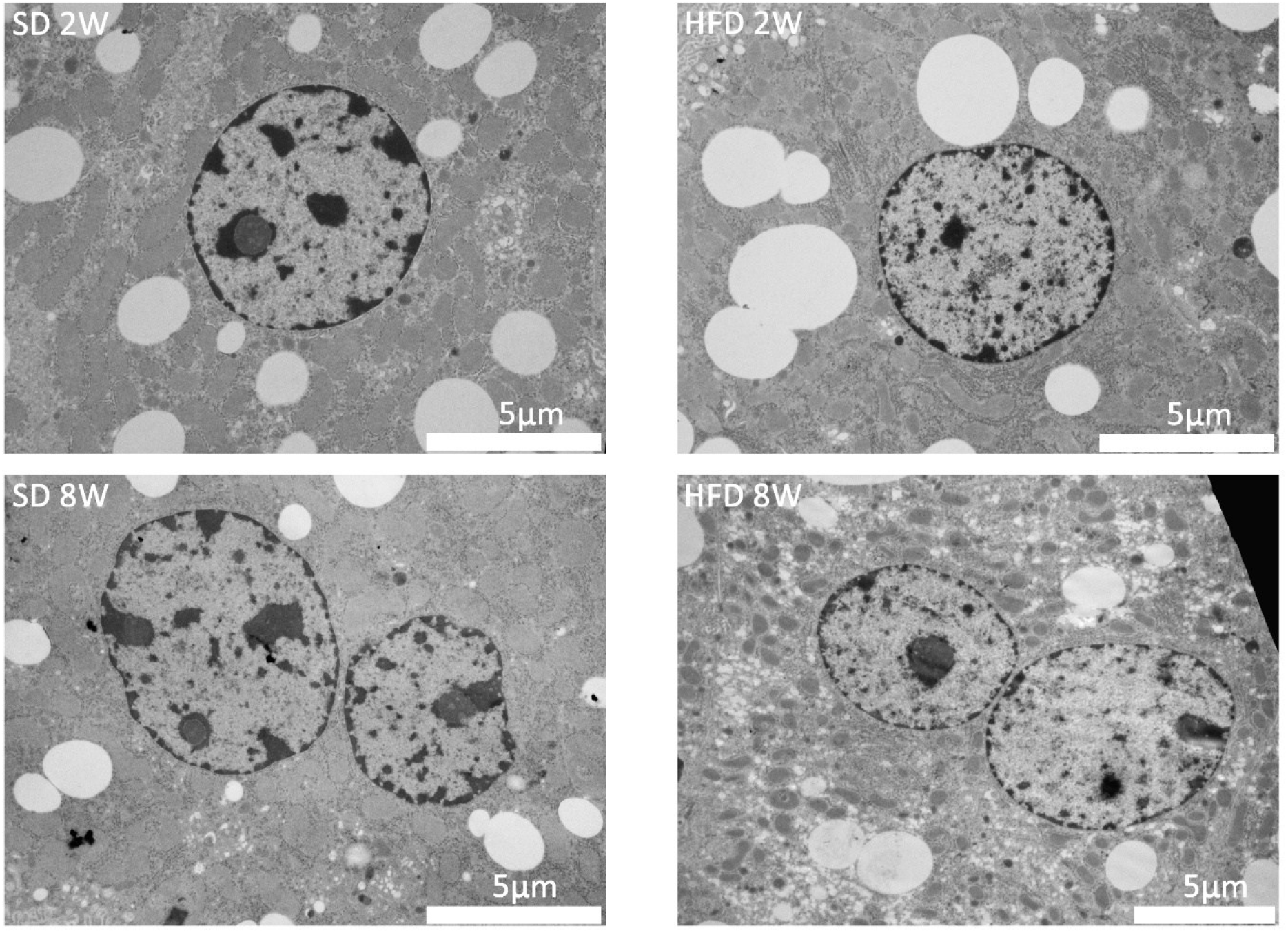
Overview of TEM ultrastructure of SD-fed vs HFD-fed mice. Representative TEM images, at 7500x magnification, of liver tissues from SD-or HFD-fed mice at 2W or 8W timepoints. Images are representative for n = 6 (SD 2W), n = 5 (HFD 2W), n = 5 (SD 8W), and n = 5 (HFD 8W) cells per condition. Bar, 5 µm.

Next, we analysed MERCS in SD-vs HFD-fed mice (Figures 6 and 7). In SD-fed mice at both time-points (2W and 8W, Figure 6A and C) mitochondria presented elongated and curved shape; most of mitochondria were covered by rough endoplasmic reticulum (RER, white arrow in Figure 6Aa’). Average transversal distance between RER membrane and OMM was (mean ± s.d.) 35.6 ± 11.9 nm at 2W of SD and 30.5 ± 10.9 nm at 8W of SD (four month of mice age) suggesting a reduction of ER-OMM distance (p < 0.0001, SD-2W vs SD-8W) (Figure 7A). A strong reduction of ER-OMM distance was found in HFD-fed mice at both 2W time-point (26.3 ± 11.3 nm) and 8W time-point (21.3 ± 7.8 nm), corresponding to a reduction by more than 25% (p < 0.0001, SD-2W vs FHD-2W; p < 0.0001, SD-8W vs HFD-8W) (Figure 7A). The reduction of ER-OMM distance was evident from a shift of most present distance range (30-40 nm) in SD-fed mice to 20-30 nm range in HFD-fed mice at 2W and 10-20 nm range in HFD-fed 8W time-point (Figure 7B). Average mitochondrial perimeter was reduced by about 37% in HFD vs SD at 2W; and by about 34% at 8W (p < 0.0001 for both comparisons). MERCS length, measured as an extension of the ER-OMM interface, was also reduced by ∼54% (p < 0.0001) and ∼38% (p = 0.0001) in HFD vs SD at 2W and 8W, respectively. In spite of the reduction of both mitochondrial perimeter and MERCS length, the percentage of mitochondrial coverage by MERCS (quantified by using a formula [% of coverage] = [MERCS length] / [MIT perimeter]) was significantly reduced in HFD-fed mice compared with SD-fed mice: by ∼40% (p < 0.0001) at 2W time-point and by ∼19% (p = 0.0079) at 8W time-point (Figure 7C). Ultrastructural analysis of TEM images at 25000x magnification confirms presence of mitochondria poorly covered by MERCS (Figure 6Bb’, white arrow) and abundant presence of free RER cisternae between mitochondria (white arrowhead), particularly visible in HFD-fed mice at 2W (Figure 6Bb’). In line with this observation, number of mitochondria per 25000x field was higher in HFD-fed mice at 2W but not at 8W (Figure 7C, MIT number). Analysis of mitochondrial area and form factors, such as circularity and diameter of Feret, confirmed smaller size and a rounder shape of mitochondria in HFD-fed mice at both time-points (Figure 7C).

**Figure 6.**
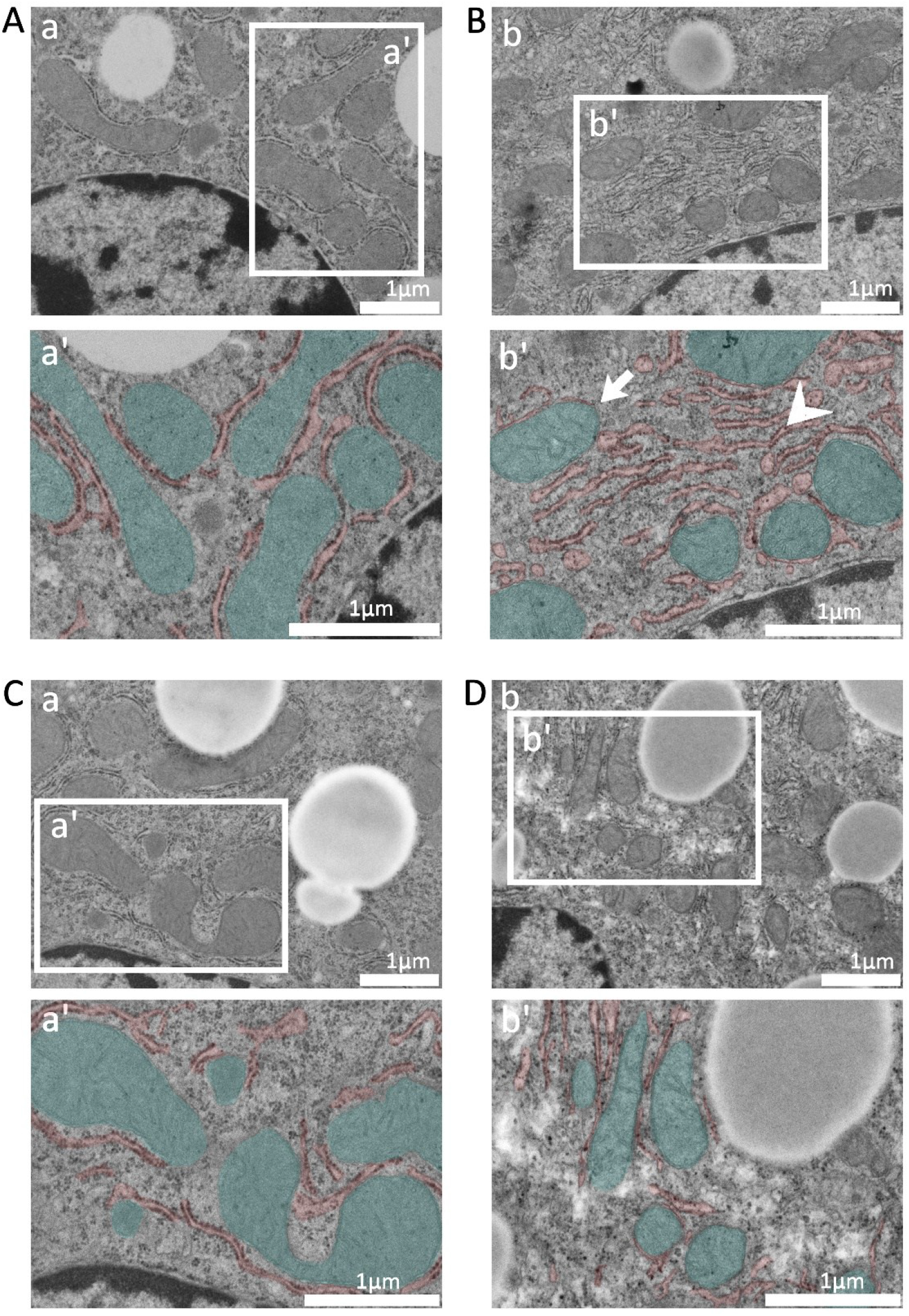
Ultrastructural analysis of MERCS. Representative images of TEM images, at 25000x magnification of liver tissues from SD-fed (**A** and **C**) or HFD-fed (**B** and **D**) mice at 2W (**A** and **B**) or 8W (**C** and **D**) time-points. White boxes delimit insets (a’ and b’) for each condition. In a’ and b’ insets, mitochondria are stained in green while ER is stained in red. Images are representative of n = 17 (SD 2W), n = 18 (HFD 2W), n = 15 (SD 8W) and n = 19 (HFD 8W) pictures per condition Bar, 1 µm.

**Figure 7.**
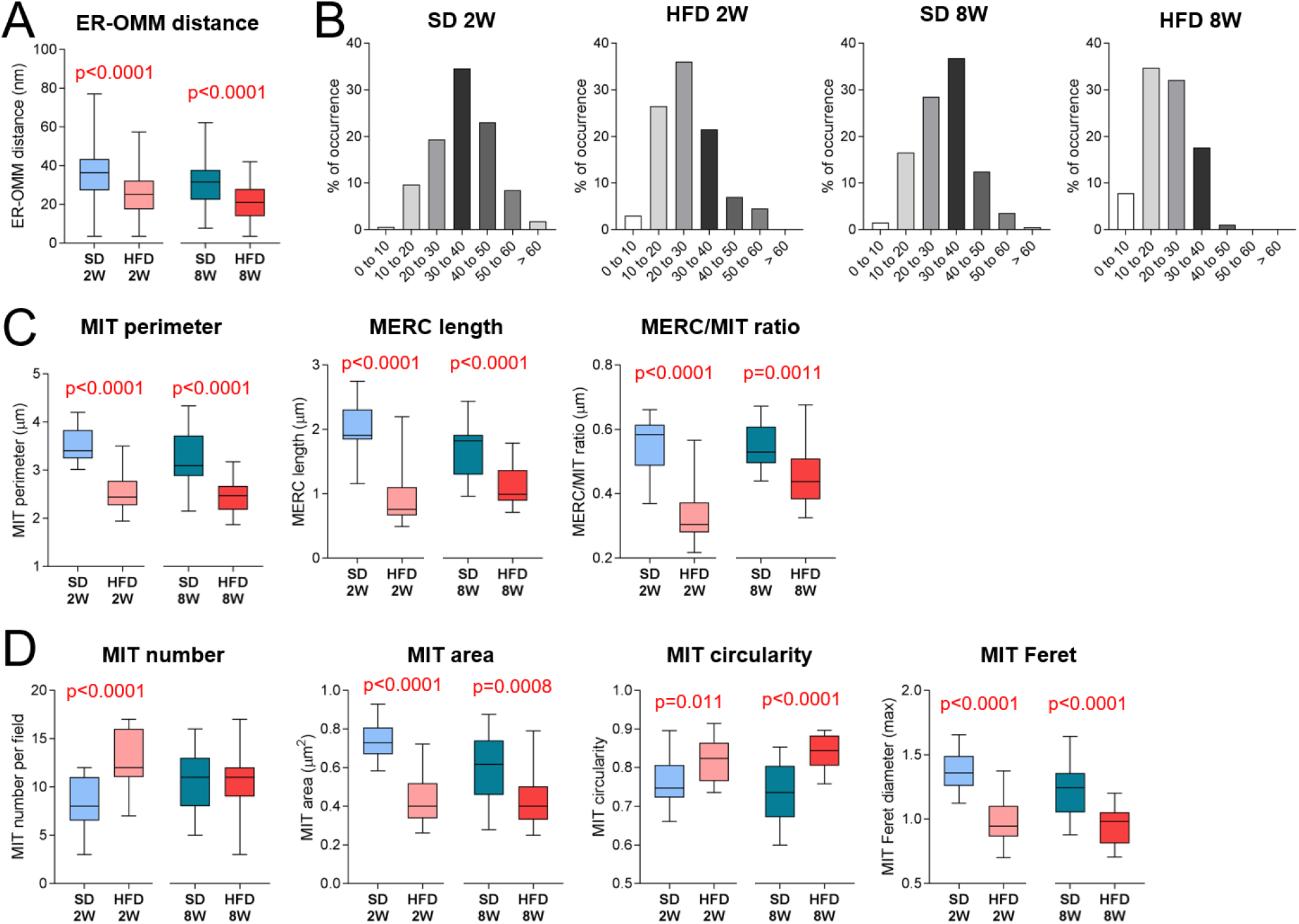
MERCS and mitochondria morphometric analysis in HFD-fed mouse liver. (A) ER-OMM distance. (B) frequencies of ER-OMM distances segmented at 10 nm intervals. (C) mitochondrial (MIT) perimeter, MERCS length and MERCS/mitochondrial perimeter ratio (MERCS/MIT ratio). (D) mitochondrial number and parameters of mitochondrial morphology: area, circularity and Feret diameter. Whisker plots in A, C and D show median, Q1-Q3 interval and maximal and minimal values. Data are collected from 140-200 mitochondria from 15-19 images per condition. In (**A** and **B**) all measurements were taken for the analysis, while in (C and D) data from microscopic fields were averaged and the averaged values were used in the analysis. P values are shown for p < 0.05.

### Spared mitochondrial function in the liver of HFD-fed mice

To assess functional alterations in mitochondria from liver of HFD-fed mice, we performed high-resolution respirometry in combination with real-time PCR expression analysis of genes induced by increased fat intake. With the exception of proton leak, increased in HFD at 2W and reduced at 8W, analysis of mitochondrial respiration by Oroboros oxygraphy did not reveal significant differences (Figure 8A, B). Although not reaching statistical significance, a trend to an increased mitochondrial respiration at 2W point was particularly evident for OXPHOS-linked activity of complex I (p = 0.055, n = 4); while the most reduced processes at 8W point were proton leak (p = 0.009), ETS respiration by complex II (p = 0.079) and by complex IV (p = 0.057, n = 3).

**Figure 8.**
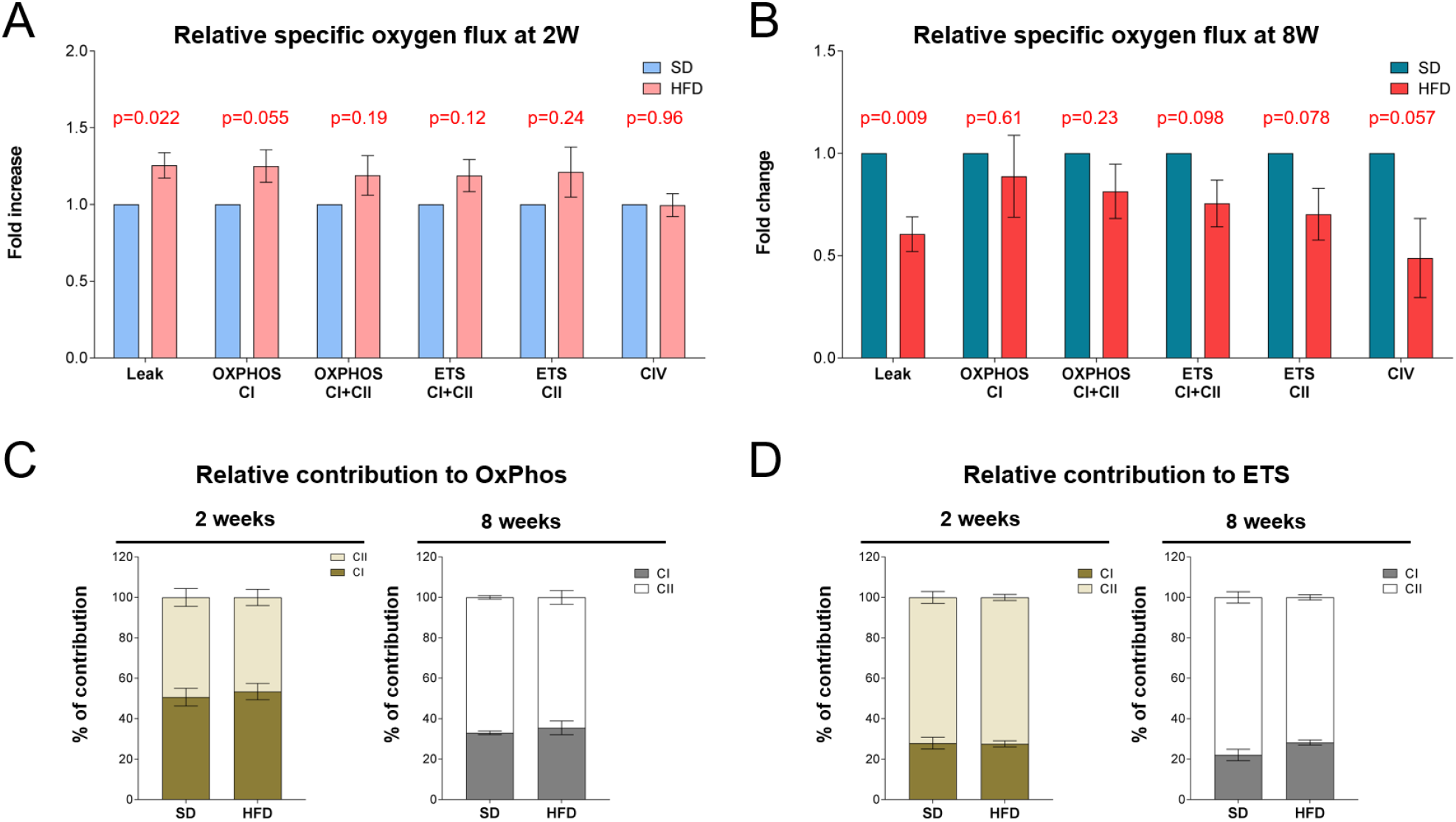
High resolution respirometry of HFD-fed mouse liver. (**A, B**) Relative oxygen consumption rate normalized to the control after 2 weeks of HFD (**A**) and 8 weeks of HFD (**B**). At the baseline, malate and pyruvate were added to induce the leakage state (Leak); ADP induces maximal oxidative phosphorylation (OXPHOS) that, in the presence of the NAD-linked substrates malate, pyruvate and glutamate, represents the OXPHOS capacity of the complex I (OXPHOS CI). The addition of succinate (S) results in the cumulative OXPHOS capacity of complexes I and II (OXPHOS CI+CII). FCCP was adjusted to obtain the maximal respiratory capacity linked to complex I and II (ETS CI+CII), while after rotenone the ETS is linked only to the complex II (ETS CII). After antimycin, complex CIV activity (CIV) was evaluated subtracting the azide-insensitive rate from the tetramethyl phenylene diamine (TMPD) + ascorbate rate. All data are expressed as specific flux, i.e., oxygen consumption normalized to the sample weight and after nonmitochondrial oxygen flux (ROX) subtraction. (**C, D**) Relative contribution of complex I and complex II in OXPHOS phase (**C**) and in ETS phase (**D**) after 2 or 8 weeks of HFD.

The differences in the electron transport system (ETS) activity at 8W time-point, if any, could be masked by the changes in relative contribution of complexes I and II to OXPHOS and ETS, e.g., [33]. However, we did not find any difference in the relative contribution of complex I or II to OXPHOS or ETS (Figure 8C, D) suggesting a good preservation of mitochondrial function.

### Alterations of lipid metabolism suggests an early activation of compensatory mechanisms in response to HFD

To get a better insight in relationships between alterations of MERCS and mitochondrial function, we performed a series of qPCR experiments assessing mRNA levels of genes, typically involved in an HFD-induced liver remodeling. First, we assessed expression of inducible regulators of lipid metabolism such as Ppar-γ, Ppar-α and Srebf1. Of these, none was significantly changed at 2W, while only Ppar-γ was upregulated at 8W time point (Figure 9A). Markers of mitochondrial and cellular stress were also assessed: Pgc-1α showed a general, although non-significant, reduction, while Cyp2e1 was up-regulated at both time-points (Figure 9B).

**Figure 9.**
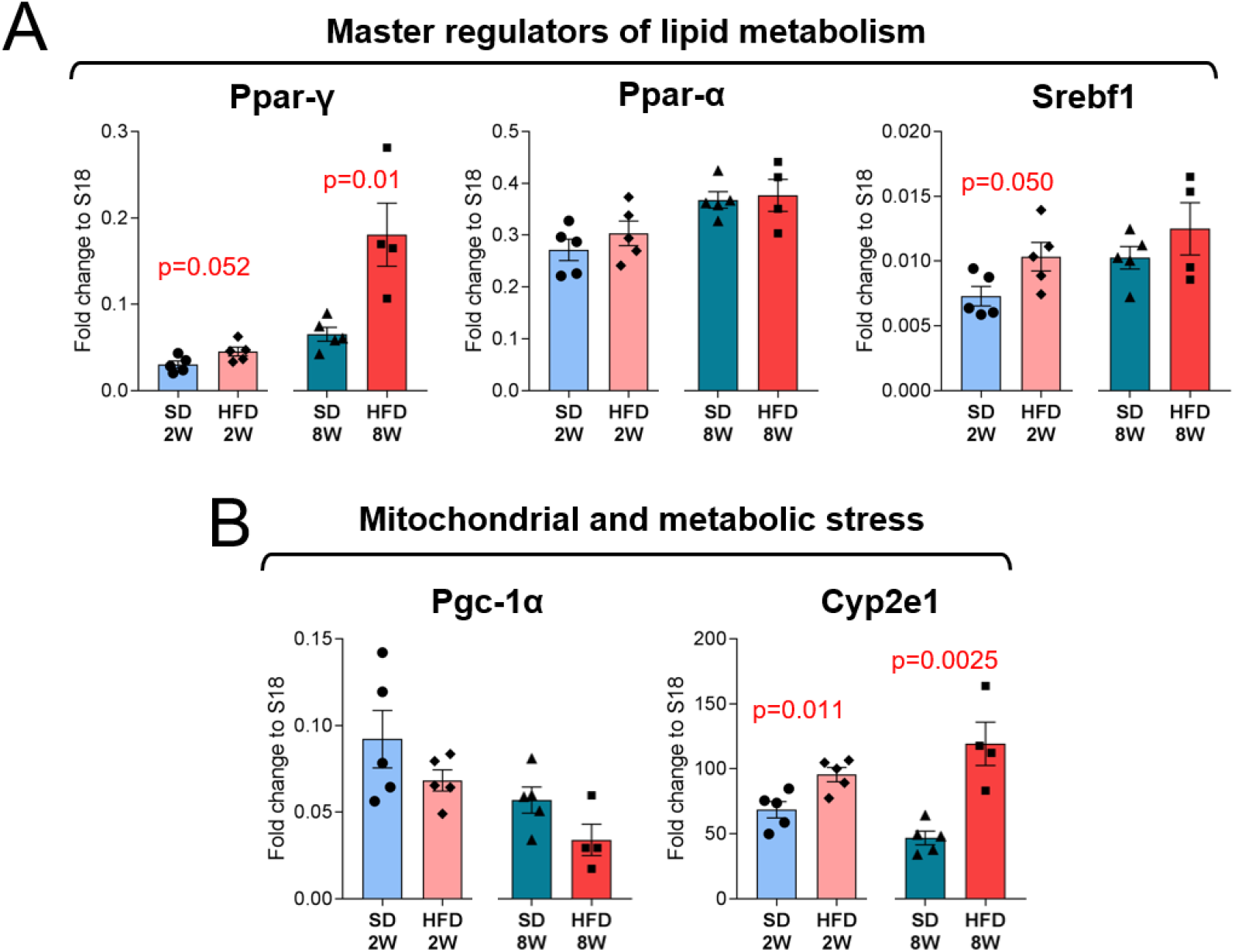
Expression of genes, involved in cellular and mitochondrial stress. Relative expression levels of inducible master regulator transcription factors Ppar-γ, Ppar-α and Srebf1 (**A**), and genes involved in cellular metabolic stress response Pgc-1α and Cyp2e1 (**B**). Dot plot graphs show mean ± s.e.m. of the gene expression fold change normalized to S18, each dot represent an independent mouse for n = 4-5 mice per condition. P values are shown for p < 0.05.

This pattern suggests an early metabolic stress, which however did not elicit a systemic transcriptional response.

Next, we assessed expression of genes involved in lipid transport and metabolism, canonically associated with increased fat intake. Genes involved in lipid transport across plasma membrane (CD36) and cytosolic lipid binding (Fabp1 and Fabp4) were largely unchanged, while Scd1 which converts saturated fatty acids into mono-unsaturated fatty acids was significantly downregulated at 2W of HFD, which was in line with previous reports [34–36] (Figure 10A). No significant changes were found at 2W time-point in the expression Ctp-1α, Acadvl and Acadl, involved in β-oxidation of fatty acids. Surprisingly, a significant reduction of Ctp-1α and Acadl was observed at 8W time-point.

**Figure 10.**
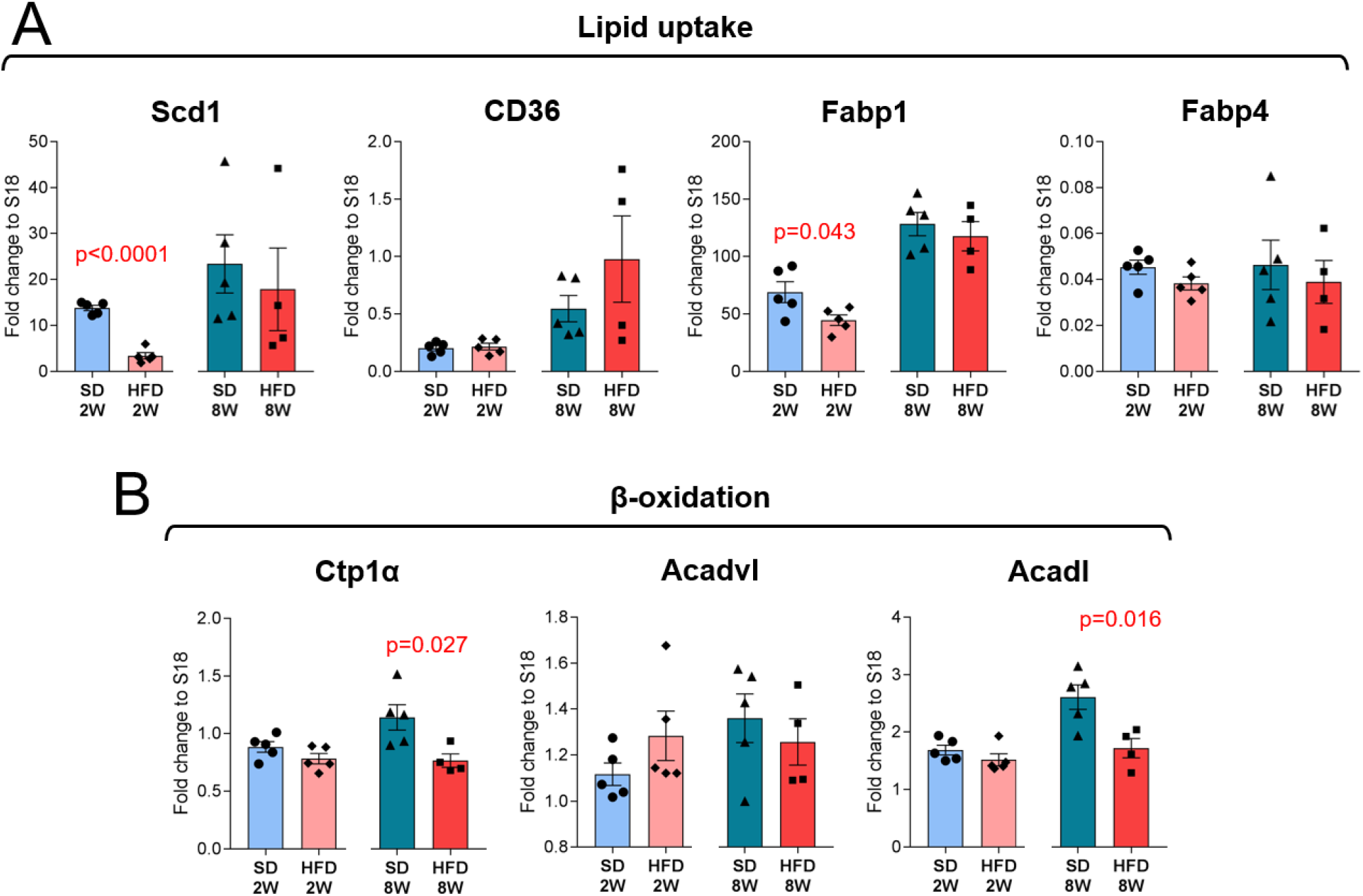
Expression of genes involved in lipid uptake and β-oxidation. Relative expression levels of genes in lipid uptake Scd1, CD36, Fabp1 and Fabp4 (**A**), and in mitochondrial β-oxidation of fatty acids Ctp1α, Acadvl and Acadl (**B**). Dot plot graphs show mean ± s.e.m. of the gene expression fold change normalized to S18, each dot represent an independent mouse for n = 4-5 mice per condition. P values are shown for p < 0.05.

Altogether, these data suggest that, at the early stage of HFD-induced liver pathology, there are very modest measurable alterations, mostly related to cellular and mitochondrial stress. At 8W time-point, only Ppar-γ was upregulated, although without systemic pathologic remodeling of lipid metabolism, which may spare mitochondrial function as compared with other reports assessing significantly longer HFD treatment [8,10,37]. Furthermore, metabolic alterations are unable to explain early disruption of MERCS in the liver of HFD-fed mice and a relative depletion of IP3R, VDAC1 and Grp75.

### Early alterations of protein homeostasis in the liver of HFD-fed mice

Recently, we suggested an association between MERCS and protein homeostasis in astrocytes, in particular with the conversion of constitutive proteasome to immunoproteasome (IP) [38–40]. Because of immunoproteasome activation is strongly implicated in HFD-induced dysproteostasis, we decided to investigate the expression of IP inducible subunits Lmp2 (β1i), Mecl-1 (β2i) and Lmp7 (β5i). As shown in Figure 11A, all three inducible IP subunits were significantly upregulated already at 2W time-point, while at 8W time-point, Mecl-1 and Lmp7 were upregulated. This finding prompted us to investigate a recently suggested axis IP → ER-stress/UPR → MERCS, postulating that ER-stress/UPR mediates MERCS disruption in conditions of lost protein homeostasis. However, we did not find activation of UPR-activated transcriptional reporters downstream of three canonical arms of ER-stress/UPR, namely Atf4 (activated by PERK), Atf6, and Xbp1s (a variant of Xbp1 transcript, alternatively spliced by IRE-1α), with an exception of Xbp1s, significantly upregulated at 8W time-point. Additionally, an ER-stress marker Herpud1 was not altered (Figure 11B). These results compellingly demonstrate that, if there is a link between IP activation and MERCS disruption, at early stage of HFD administration, it does not require activation of UPR.

**Figure 11.**
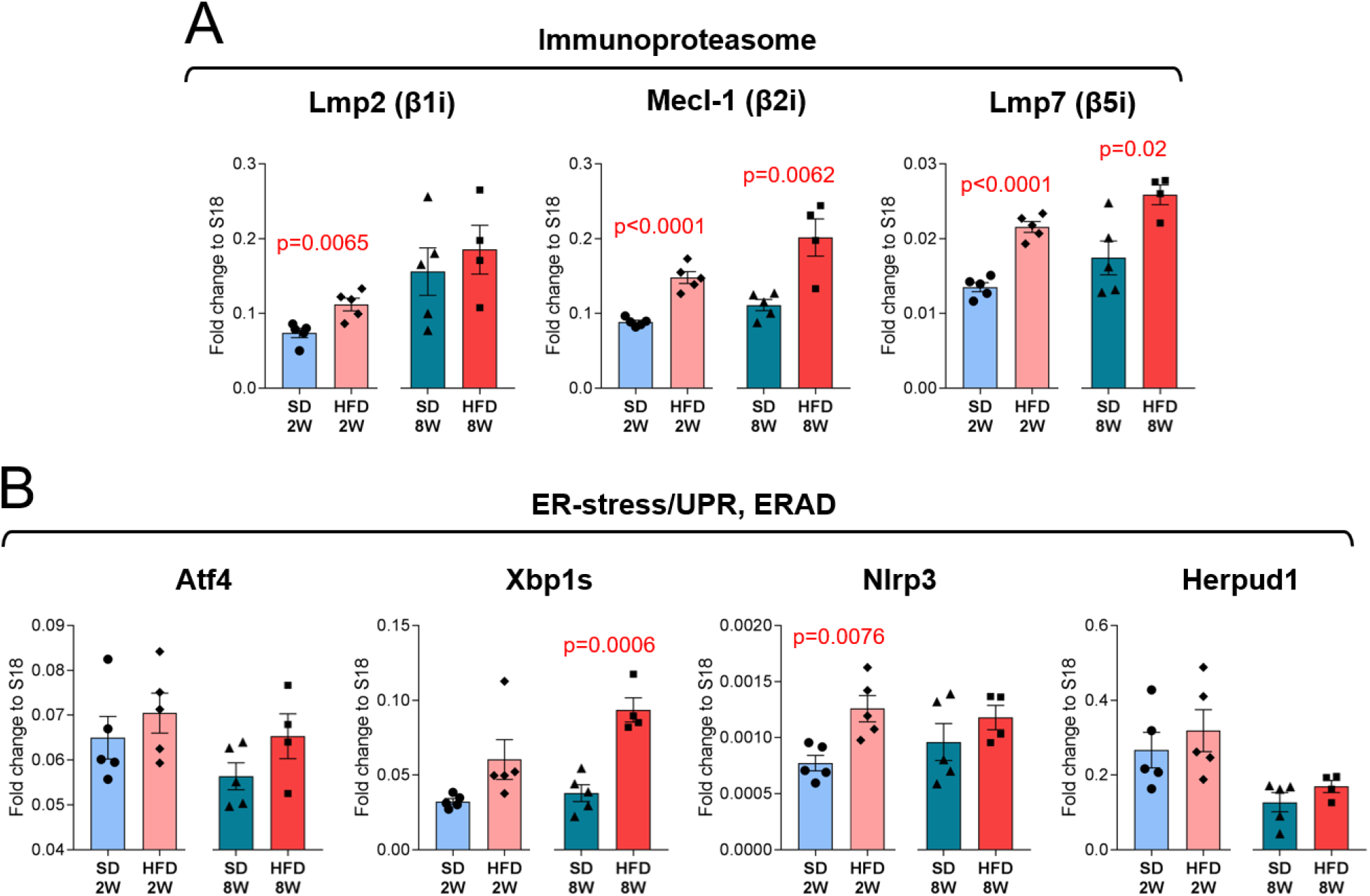
Immunoproteasome and ER-stress/UPR. Relative expression levels of β1i, β2i and β5i inducible immunoproteasome subunits (**A**), and genes, typically induced downstream of ER stress (**B**). Dot plot graphs show mean ± s.e.m. of the gene expression fold change normalized to S18, each dot represent an independent mouse for n = 4-5 mice per condition. P values are shown for p < 0.05.

## DISCUSSION

The specific aim of this study was to correlate the expression of IP3R, VDAC1 and Grp75 in MERCS with their ultrastructural alterations, specifically physical parameters of this interaction such as transversal ER-OMM distance and mitochondrial coverage with MERCS and distribution, in SD vs HFD-fed mice at two time points of 2W and 8W. These were correlated with mitochondrial function and expression of genes associated with high fat load. Main finding of this study include: i) early reduction of the expression of IP3R at a posttranscriptional level; ii) early alterations of mitochondrial morphology and the reduction of mitochondria coverage by MERCS as well as a reduction of transversal ER-mitochondria distance in remaining MERCS; and iii) depletion of IP3R, VDAC1/3 and Grp75 in MERCS from MERCS at an advanced stage (8W).

Regarding the alterations of MERCS in models of obesity and increased food intake, somewhat controversial results were published. However, analysis of the reports suggests that early alterations in MERCS during HFD administration (1-2 weeks) are consistent with the reduction of ER-mitochondrial communication in terms of mitochondrial coverage by MERCS, and the reduction of hepatocytic Ca^2+^ signals and/or sensitivity to stimulation [10,41]. Present work supports this observation, we found both downregulation of IP3R in total homogenates and a reduction of ER-mitochondrial communication. The controversies are mostly related to long-term HFD treatment, lasting up to 12-16 weeks. The reasons accounting for the discrepancies might include different housing conditions, diet formulations and administration protocols applied for longer period. Besides these, experimentally intrinsic variables, variations in the parameters of evaluation, inclusion criteria and algorithms of quantification of morphometric parameters of MERCS add to divergent results and conclusions [8,10,42]. Nevertheless, genetic models of obesity and diabetes consistently show increased expression of IP3Rs and Ca^2+^ signals [8,43], suggesting that upstream mechanistic aspects are important factors and should be taken in consideration while evaluating MERCS alterations.

Another observation of this work, which deserves a brief discussion, is that at 2W of HFD, when strong reduction of MERCS length and a downregulation of IP3R was observed, mitochondrial function was largely spared with modest alterations of lipid metabolism, as judged by the expression of the related inducible enzymes. It is widely accepted that ER-mitochondrial Ca^2+^ flux and a low affinity mitochondrial Ca^2+^ uptake drive bioenergetic metabolism in the mitochondrial matrix [19]. Therefore, one could expect that a reduction of ER-mitochondrial communication together with downregulation of IP3Rs would have strong negative impact on mitochondrial energetic metabolism. However, alongside the reduction of MERCS coverage and length, we found a significant shortening of the transversal ER mitochondrial distance, specifically from 35.6 ± 11.9 nm to 26.3 ± 11.3 nm at 2W time-point and from 30.5 ± 10.9 nm to 21.3 ± 7.8 nm at 8W of HFD. Notably, the reduction in MERCS extension occurred in parallel with the shortening of ER-mitochondrial distance, suggesting a structural reorganization of ER-mitochondria contacts rather than a simple loss of interaction. These structural changes, together with early IP3R down-regulation, suggest a temporal sequence in which MERCS remodeling precedes the depletion of the IP3R-Grp75-VDAC complex and occurs before detectable mitochondrial dysfunction. Recently, we reported that a distance of about 20 nm between ER and OMM is optimal for ER-mitochondrial Ca^2+^ transfer and uptake, while at a distances ≥ 30 nm the efficiency of ER-mitochondrial Ca2+ flux is dramatically reduced [28]. Moreover, stabilization of ER-mitochondrial interaction at about 20 nm rescues mitochondrial Ca^2+^ uptake, bioenergetics and cellular proteostasis in experimental and pathological conditions including Parkinson’s and Alzheimer’s disease astrocytes [28,39]. Although this mechanism(s) might explain how a relative reduction of ER-mitochondrial distance could enhance ER-mitochondrial Ca^2+^ flux and, in some extent, compensate for the reduction of ER-mitochondrial interface, this remains a speculation; many other factors need to be considered, including change in substrate usage, alterations of membrane lipid composition and adaptive mechanisms. Certainly, direct assessment of Ca^2+^ flux between ER and mitochondria would help to interpret these data. Unfortunately, due to technical limitations and legal constrains, we were not able to assess Ca^2+^ flux and mitochondrial Ca^2+^ uptake; we admit that this constitutes a strong limitation of this work. Nevertheless, available literature suggests that, in similar conditions both cytosolic and mitochondrial Ca^2+^ signals are reduced [10,41].

Regarding the cause-effect between MERCS and cell dysfunction, canonical view postulates that HFD first, induced loss of protein homeostasis resulting in ER-stress with activation of UPR response. This, in turn, results in alterations of ER-mitochondrial communication with consequences for lipid and energetic metabolism [8,44–49]. However, recent reports and the present data challenge this scenario, suggesting that the loss of MERCS stability triggers dysregulation of protein homeostasis [10,50]. In line with this “MERCS - first” hypothesis are also our recent reports showing that stabilization of MERCS at a defined distance reverted IP activation and rescued autophagic flux in hippocampal astrocytes from a mouse model of Alzheimer’s disease [39]. Further experiments, with carefully defined and harmonized experimental conditions and analytical processing will be necessary to clarify the role of MERCS alterations in liver dysfunction in metabolic diseases and obesity.

Altogether, alterations of mitochondria and MERCS, and the reduced expression of IP3R in hepatocytes, represent an early event during increased caloric and fat intake, occurring well before a gain of weight and overt obesity or development of glucose intolerance. Our results provide further support in suggesting that MERCS alterations may represent a triggering event for metabolic dysfunction and consequent complications, promoting MERCS as a potential target for future therapeutic approaches to counteract the widespread burden of metabolic diseases, obesity and diabetes. Because of NAFLD and NASH represent risk factors for age-related dementias, including Alzheimer’s disease [4–6], we suggest that targeting MERCS in the liver at early stages of metabolic dysfunction may also represent a preventive measure towards healthy aging.

## MATERIALS AND METHODS

### Mice and diet administration

The same mice cohort used in [29] was used in this study. Male C57BL/6J mice were used in this study. Animals were housed 3–5 per cage, with free access to food and water (ad libitum). They were maintained in a temperature-controlled environment (22–24 °C) with a 12 h light/12 h dark cycle in high-efficiency particulate air (HEPA)-filtered Thoren units (Thoren Caging Systems) at the animal facility of the University of Piemonte Orientale. All procedures involving animal care and handling complied with the Italian legislation on animal welfare (D.L. 26/2014) and the European Directive (2010/63/EU). The experimental protocol was approved by the Organismo Preposto al Benessere Animale (OPBA) of the University of Piemonte Orientale, Novara, Italy (DB064.61).

Thirty-eight mice began receiving a low-fat diet (SD; 13% kcal from fat, 67% kcal from carbohydrates, and 20% kcal from proteins; Laboratori Piccioni, Milan, Italy) starting at four weeks of age, replacing the standard chow diet normally provided by the animal facility in order to acclimate the animals to a refined diet [51]. After one week (five weeks of age), mice were randomly assigned to four groups. Two groups continued to receive the low-fat diet (control groups) for either eight weeks (n = 9) or two weeks (n = 9). The remaining two groups were switched to a high-fat diet (HFD; 60% kcal from fat, 21% kcal from carbohydrates, and 19% kcal from proteins; Laboratori Piccioni) for eight weeks (n = 10) or two weeks (n = 10). Body weight, as well as food, water, and caloric intake, were monitored weekly. Glycemia was evaluated in mice fasted for 18h before the beginning of the experiment and after 2 and 8 weeks of diet regimen. At the end of the dietary interventions, mice were euthanized and liver samples were collected. Tissues were snap-frozen at −80 °C for subsequent biomolecular analyses or processed for TEM analysis as described below.

### MERCS isolation

Isolation of MERCs from mouse liver was performed based on a modified protocol described by Wieckowski et al [30]. Single MERCs preparation was obtained from two liver lobules collected from two mice belonging to the same control or experimental group. Three and four independent preparations were performed for 2W and 8W time-point, respectively. As the first step, liver tissue fragments were washed out from the blood performing one wash in IB_liver_-1 buffer, which was subsequently substituted with Starting Buffer. Liver fragmentation was performed in IB_liver_-1 buffer supplemented with phosphatase inhibitor cocktail (Cell Signaling Technology, Cat. 5870) and protease inhibitor cocktail (PIC, Millipore, Cat. 539133). Tissue was finely intersected with scissors and then homogenized using a glass Teflon homogenizer, tearing tissue manually with Teflon pestle performing 25 strokes, yielding total homogenate fraction (HOM).

The homogenate was transferred to the Eppendorf tubes and centrifuged at 750 x *g* for 5 min at 4 °C. Obtained supernatant was collected, while remaining pellet was resuspended in 200 µl IB_liver_-1 complemented with protease and phosphatase inhibitor cocktails, homogenized and centrifuged again at 750 x *g* for 5 min at 4 °C, subsequently the supernatant was collected and pellet discarded. Collected supernatant was centrifuged at 9000 x *g* for 5 min at 4 °C, further resulting pellet was gently resuspended in 1 ml of IB_liver_-2 buffer with addition of protease and phosphatase inhibitor cocktails and centrifuged at 10,000 x *g* for 10 min at 4 °C. Obtained pellet representing crude mitochondrial fraction was resuspended in 950 µl MRB buffer containing protease and phosphatase inhibitor cocktails.

Percoll gradient was prepared, adding 16 ml Percoll medium to the ultracentrifuge tube, subsequently crude mitochondria fraction was carefully loaded on the Percoll medium and additional 3 ml MRB were delicately layered on top of the gradient. The samples were ultracentrifuged at 100 000 x *g* for 30 min at 4 °C, using an Eppendorf CR30NX ultracentrifuge equipped with an R25ST rotor. The MERCS fraction was collected and diluted with 20 ml MRB and further ultracentrifuged at 100 000 x *g* for 1 h at 4 °C. In the final step, obtained pellet representing purified MERCS fraction was resuspended in MRB and mixed in ratio 1:1 with lysis buffer (50 mM Tris-HCl pH 7.4, 0.5% SDS, 5 mM EDTA) supplemented with protease and phosphatase inhibitor cocktails, and stored at −20 °C.

Isolation of MERCS was performed from liver tissue in parallel on the same day for both SD and HFD experimental groups, but in different days for 2W and 8W time-points.

### Western blot

Protein quantification of obtained fractions was performed using QuantifiPRo BSA Assay Kit (Sigma, Cat. SLBF3463). Depending on the relative abundance of protein of interest, 20-40 µg proteins were combined with appropriate volume of Laemmli Sample Buffer 4X (Bio-Rad) and boiled. Samples were loaded on 6% or 12% polyacrylamide-sodium dodecyl sulfate gel for SDS-PAGE. Proteins were transferred onto nitrocellulose membrane, using Mini Transfer Packs or Midi Transfer Packs, with TransBlot® Turbo ™ (Bio-Rad), according to the manufacturer’s instructions. The nitrocellulose membranes were blocked in 5% skim milk (Sigma, Cat. 70166) for 1h at room temperature. In the following step, the membranes were incubated with indicated primary antibody overnight at 4 °C. For the normalization of loading of proteins, anti-β-Actin was used. Details of primary antibodies is provided in Table 2. Goat anti-mouse IgG (H+L) horseradish peroxidase-conjugated (BioRad, 1:5000; Cat. 170-6516) and Goat anti-rabbit IgG (H+L) horseradish peroxidase-conjugated secondary antibodies (Bio-Rad, 1:5000; Cat. 170-6515) were used. Detection was carried out using SuperSignalTM West Pico/ femto PLUS Chemiluminescent Substrate (Thermo Scientific, Cat. 34578) and was detected using ChemiDocTM Imaging System (Bio-Rad).

**Table 2.** List of primary antibodies used in this study.

| Primary Antibodies | Animal specificity | Dilution | Cat.N. | Supplier |
| --- | --- | --- | --- | --- |
| Anti- IP3R1 | Rabbit | 1:500 | AB108517 | Abcam |
| Anti-VDAC 1/3 | Mouse | 1:1000 | AB14734 | Abcam |
| Anti-GRP75 | Rabbit | 1:1000 | 14887-1-AP | Proteintech |
| Anti-Mitofusin 1 | Rabbit | 1:200 | SC-50330 | Santa Cruz |
| Anti β -Actin | Mouse | 1:5000 | A1978 | Sigma-Aldrich |

### 2.4. Real-time PCR

Total mRNA was extracted from 10-20 mg of liver using TRIzol Lysis Reagent (Invitrogen, Cat. 15596026) according to manufacturer’s instruction. First strand of cDNA was synthesized from 1.2 µg of total RNA using SensiFAST cDNA synthesis kit (Meridian Bioscience, UK, Cat. BIO-65054). Real-Time PCR was performed using iTaq qPCR master mix according to manufacturer’s instructions (Bio-Rad, Cat. 1725124) on a SFX Opus 96 Real-time system (Bio-Rad). To normalize raw real time PCR data, S18 ribosomal subunit was used. Data are expressed as fold change compared with SD-fed mice. Sequences of oligonucleotide primers are present in Table 3.

**Table 3.**
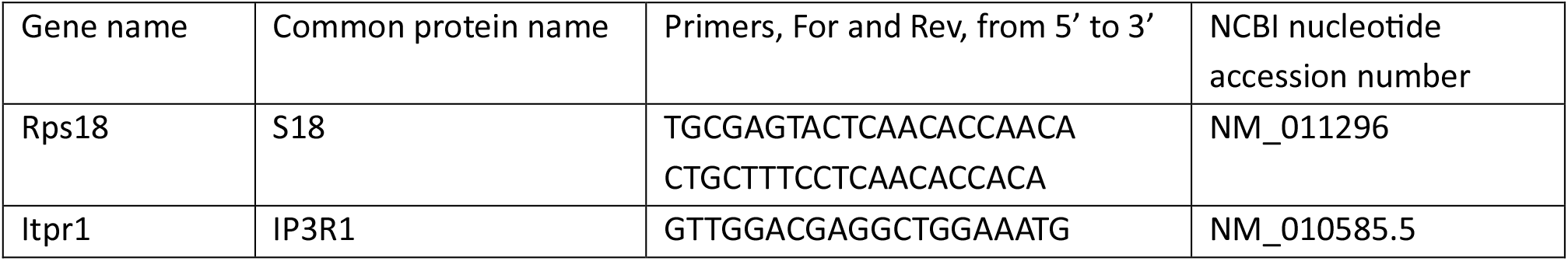
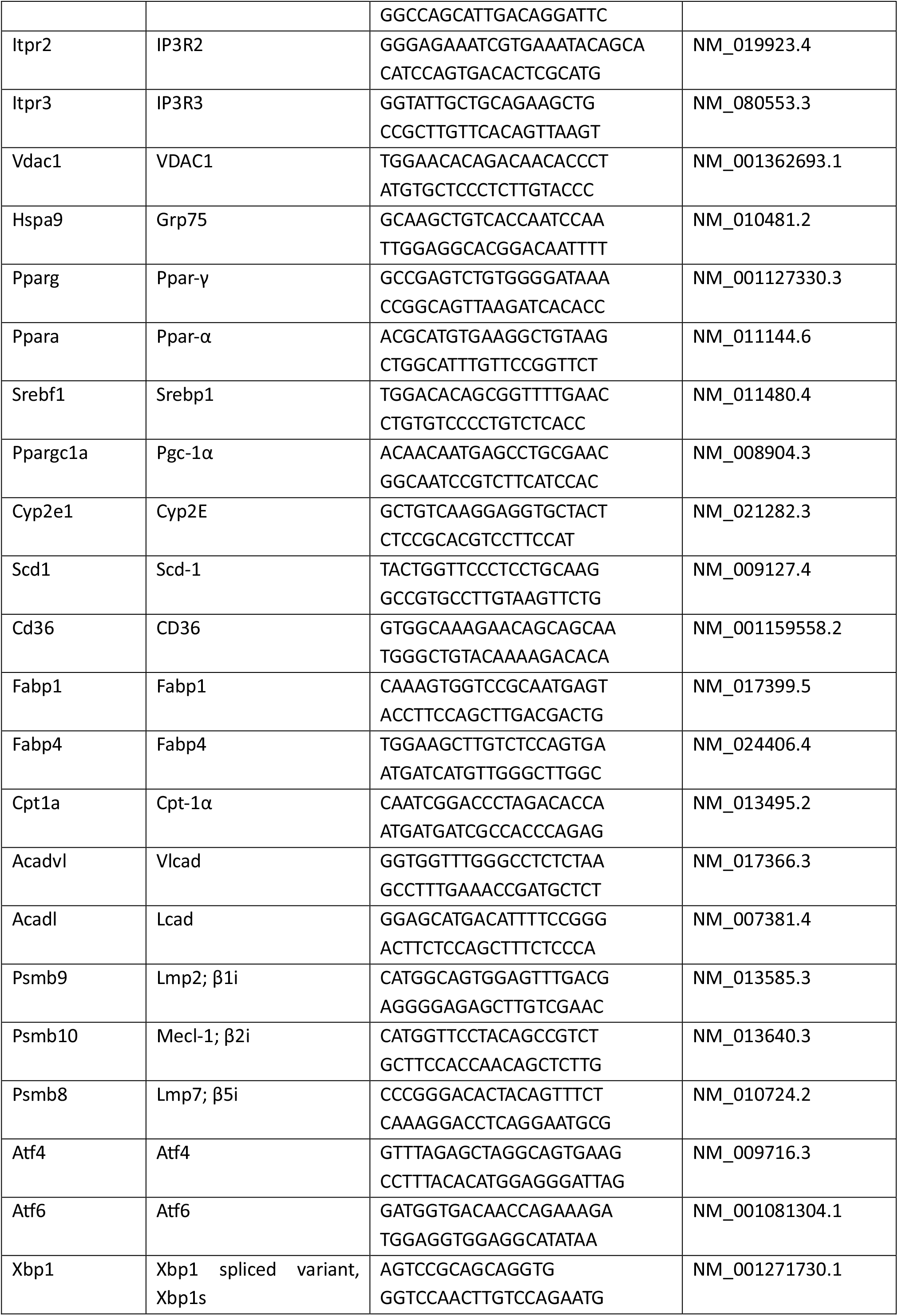
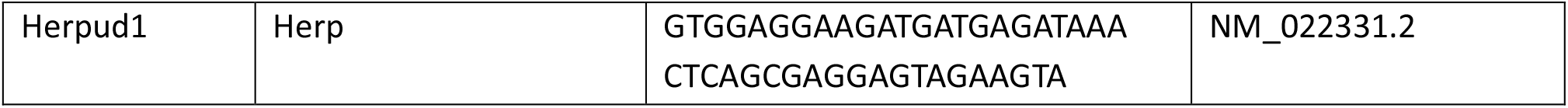
Oligonucleotide primers used for real-time PCR.

### Transmission electron microscopy

For transmission electron microscopy (TEM) analysis, liver biopsies of about 1 mm^3^ were fixed in Karnovsky fixative for 2 h at room temperature. Samples were then rinsed, post-fixed in 1% aqueous OsO_4_ for 2 h at room temperature and rinsed in H_2_O. Tissues were dehydrated in a graded acetone scale and then embedded in epoxy resin (Electron Microscopy Sciences, EM-bed812). Ultrathin sections (60–80 nm) were cut on a Reichert OM-U3 ultramicrotome, collected on nickel grids, and then stained with uranyl acetate and lead citrate. The specimens were observed with a JEM 1200 EX II (JEOL, Peabody, MA, USA) electron microscope operating at 100 kV and equipped with a MegaView G2 CCD camera (Olympus OSIS, Tokyo, Japan) [52,53]. Images were analyzed via ImageJ 1.54F.

For the ER-mitochondrial distance/interface analysis, the regions of juxtaposition between ER membrane and OMM were identified with a gap from 0 to 80 nm. These regions were considered as MERCS and the length of their extension was quantified as “MERCS length” or “length of the ER-mitochondria interaction”. Transversal distance between ER and OMM was quantified manually using ImageJ “measure” tool. Within a single contact site with an extended interface, the transversal distances were measured along the interface with ∼20 nm intervals. A single contact of the same distance, defined by the continuous interface between ER and OMM, was considered as one point for statistical analysis. Mitochondrial morphology was assessed by manually tracing OMM perimeter and morphological parameters were quantified using ImageJ “measure” tool.

### Liver homogenization and Oroboros high resolution respirometry

High resolution Oroboros respirometry analysis was performed on three or four mice per time-points 2W and 8W, respectively. The entire liver was excised, and a small piece of no more than approximately 50 mg was removed, washed in saline (0.9% NaCl) solution, blotted on filter paper and finally weighted. After these operations, for short term storage the excised tissue was kept at 4°C in 1 ml of BIOPS solution [50 mM MES, 20 mM taurine, 0.5 mM dithiothreitol, 6.56 mM MgCl2, 5.77 Mm Na2ATP, 15 mM Na2Phosphocreatine, 20 mM imidazole, 10 mM Ca-EGTA buffer (2.77 mM CaK2EGTA + 7.23 mM K2EGTA; 0.1 μM free calcium) pH 7.1 adjusted with 5 N KOH]. To obtain liver lysate, the sample was transferred in the pestle homogenizer containing MiR05 buffer (0.5 mM EGTA, 3 mM MgCl2, 60 mM K-lactobionate, 20 mM taurine, 10 mM KH2PO4, 20 mM HEPES, 110 mM sucrose, and 1g/l BSA, pH 7.1) at a volume adjusted to achieve a ratio of 1 mg of tissue per 10 μL. The homogenizer was used at 600 rpm for six times up and down. Subsequently, 2 mg of tissue (20 μl of homogenate) was placed into the respirometer chambers containing 2 ml of the respiration medium MiR05.

Mitochondrial respiratory capacity was measured with a high-resolution respirometer (Oxygraph-2k, Oroboros Instruments, Innsbruck, AT) and the data were recorded using Datlab software (Oroboros Instruments, Innsbruck, AT). For this type of analysis, the respiration was assessed by a substrate-uncoupler-inhibitor-titration (SUIT) protocol SUIT-008 O2 ce-pce D025, as recommended by the manufacturer of the Oroboros instrument [54,55] at 37°C with the following sequential injections: 1) 1 mM malate, 5 mM pyruvate to determine non-phosphorylating LEAK respiration supported by complex I-linked substrates; 2) 5 mM ADP to achieve maximal phosphorylating respiration from electron input through complex I (OXPHOS CI capacity); 3) 10 µM cytochrome c to assess the integrity of the outer mitochondrial membrane; 4) 10 mM glutamate (OXPHOS CI capacity); 5) 10 mM succinate to saturate complex II and achieve maximal convergent electron flux through complex I and II (CI and CII OXPHOS capacity); 6) 0.5 µM carbonylcyanide m-cholorophenyl hydrazone (FCCP) to assess complex I and II linked maximal capacity (CI and CII ETS capacity); 7) 0.5 µM rotenone to inhibit complex I (ETS CII capacity); and 8) 2.5 M antimycin A to inhibit complex III and obtain residual oxygen consumption (ROX). Complex IV activity was evaluated after antimycin A CIII inhibition by ascorbate (0.5 mM) and TMPD (N, N, N’, N’-tetramethyl-p-phenylenediamine; 2 mM) injection. At the end 200 mM Azide was titered to inhibit CIV. Oxygen fluxes of the different respiratory states were corrected by subtracting residual oxygen consumption following antimycin A treatment (ROX) and normalized to the tissue weight (2 mg).

### Statistical analysis

Statistical analysis and graphs were prepared using GraphPad PRISM software versions 6-10. For statistical analysis of body weight, Two-way ANOVA with Sidak post-hoc test was used. 2W samples were processed independently of 8W samples and were analysed by two-tail unpaired Student’s t-test. For three or more conditions, one-way ANOVA was used followed by Tukey post-hoc test. Differences were considered significant with p < 0.05. Western blot results are expressed as bar-plots as mean ± s.e.m. MERCS and mitochondria morphometric data are presented by whisker plots expressing median, interquartile range (Q1 – Q3), minimal and maximal measured values. Oroboros oxygraphy results are expressed as bar-plots expressing mean ± s.e.m. qPCR results are presented by dot plot graphs showing mean ± s.e.m. of number of mice run in triplicate.

## Author Contributions

Conceptualization, L.T., F.C. D.G. L.D.; methodology, N.C., S.R., M.B.; validation, J.M, G.C., G.D.; investigation, J.M., G.C., C.C., E.P., E.M., S.R., L.T., F.C., G.D.; resources, G.P., M.B.; data curation, J.M., G.C., F.C., G.D., D.L.; writing—original draft preparation, D.L.; writing—review and editing, J.M., G.C., L.T., F.C., G.D., D.L.; supervision, L.T., F.C., G.D., D.L.; project administration, F.C., D.L.; funding acquisition, F.C., D.L. All authors have read and agreed to the published version of the manuscript.

## Acknowledgments

The authors acknowledge Imaging and Metabolic Facilities of Interdepartmental Center for Autoimmune and Allergic Diseases (CAAD), Università del Piemonte Orientale, Novara, 28100, Italy; Dr. Massimo Boiocchi, Transmission Electron Microscopy Facility, Centro Grandi Strumenti, Università di Pavia, Pavia, 27100 Italy.

## Conflicts of Interest

The authors declare no conflicts of interest.

## Notes

### Competing Interest Statement

The authors have declared no competing interest.

